# Cobalt and nickel ion synergy promotes gene duplication amplification and enables stable metal resistance in *Shewanella oneidensis*

**DOI:** 10.1101/2025.08.08.669375

**Authors:** Alessandra G. Gavin, Stephanie L. Mitchell, Erin E. Carlson

## Abstract

Microbial species are integral in environmental homeostasis and have thus developed diverse strategies to adapt, survive, and proliferate under exogenous stress. The growing demand for nanoscale battery materials has led to increasing environmental concentrations of nanomaterials and their constituent metal ions. Previous work has shown that the gram-negative bacterium *Shewanella oneidensis* is able to rapidly evolve resistance to one such nanomaterial, lithiated nickel manganese cobalt oxide (NMC). Yet, the specific stimuli that trigger resistance evolution and the ensuing genomic changes were previously unknown. Here, we demonstrate that the combination of cobalt and nickel ions released from NMC trigger a gene duplication amplification (GDA) event that enables stable nanomaterial and metal resistance. Amplification copy number is highly dynamic over the period of study, yet amplification events persist over prolonged recovery in the absence of metal stress. Growth rate comparisons reveal no physiological cost associated with high copy GDA. The stability of this genomic mutation combined with the lack of observable fitness cost distinguishes this genomic perturbation from previously reported GDA events. Lastly, the observed GDA event was unique to the combination of cobalt and nickel, which implies that the intracellular targets of these metal ions have a specific interaction that yield resistance. Ultimately, we report a dynamic and specific GDA event that allows *S. oneidensis* to survive metal stress. This work not only illuminates the broader ecological consequences associated with introducing nanomaterials and metals into the environment but also provides insight into the larger scope of bacterial resistance mechanisms.

**Importance:** Studying mechanisms of microbial adaptation and resistance evolution is critical for understanding the role of bacteria in the environment and developing new strategies to combat antimicrobial resistance to modern therapeutics. Technological innovation has often led to the introduction of novel environmental stressors that impart pressure on microbes to evolve resistance. This may ultimately lead to novel resistance genes or gene cassettes in the environmental gene pool, which could further promote antimicrobial resistance in pathogenic organisms. In this study, we highlight the importance of identifying GDA events as a mechanism of bacterial resistance to environmental toxins. We demonstrate that amplification events can exist within a population in the absence of selection pressure and without a clear fitness cost. Identifying specific stimuli that trigger these events will help understand factors that accelerate bacterial resistance evolution and have the potential to disrupt the balance of the ecosystem.

## 1. Introduction

Bacterial species are integral components of the environmental microbiome and are known to rapidly adapt and evolve under selective pressure.^1^ This phenotypic plasticity allows bacteria to temporarily survive in diverse and hostile environments and creates the opportunity for cells to acquire beneficial genomic mutations that promote sustained survival against specific stressors.^2–5^ Yet, the mutations that facilitate bacteria survival often come at the cost of fitness and thus may impact the composition of microbial communities.^6, 7^ The cost is often worth paying as these adaptive strategies have enabled bacteria to develop resistance to a diverse array of antibiotics and other toxins, including engineered nanomaterials (ENMs) and metal ions.^8^ The use of ENMs in consumer goods has increased dramatically over the past two decades, and has resulted in a detectable increase in the prevalence of ENMs and associated raw materials in waste streams and the environment.^9^ Although many of these materials are ecologically benign, some ENMs and associated metal ions have demonstrated toxicity to a variety of organisms.^10, 11^

Nanoscale lithiated nickel manganese cobalt oxide (NMC) is a commonly used cathode material in electric vehicle batteries and research has established its toxicity towards bacterial species. NMC toxicity stems from a combination of reactive oxygen species (ROS) and cobalt and nickel metal ions released from NMC in an aqueous environment.^12^ Furthermore, challenges associated with developing large-scale recycling infrastructure makes environmental release of NMC inevitable.^13–15^ Evaluating bacterial response to ENM stress is not only critical for understanding the ecological impact of environmental toxins, but also furthers our understanding of fundamental mechanisms of bacterial adaptation and evolution.

*Shewanella oneidensis* is a ubiquitous, soil-dwelling bacteria known for its role in environmental metal cycling. Given the prevalence of *S. oneidensis* in diverse aquatic environments, it is likely that *S. oneidensis* will be present in nanomaterial-contaminated environments.^16, 17^ We have previously demonstrated that chronic exposure to NMC or associated metal ions causes rapid and stable resistance in *S. oneidensis.*^18^ Understanding the physiological changes associated with metal and NMC resistance is critical due to the role of *S. oneidensis* environmental metal cycling and maintaining environmental homeostasis.^19, 20^ By understanding how exogenous stressors fundamentally change *S. oneidensis*, we can more clearly assess the environmental consequences associated with metal ions and nanomaterials.

When bacteria first encounter exogenous stress, cells activate a general stress response that increases transcriptional regulation of protective genes.^21, 22^ This initial regulatory response can have a dramatic effect on bacterial physiology and is often accompanied by reduced growth rate to enable short-term survival, but does not promote sustained growth under stress or resistance evolution.^23^ ^24, 25^ Notably, when cells are removed from direct stress, they lose these specific stress adaptations and return to being phenotypically and genotypically identical to wild-type (WT) cells.^26^ Under chronic stress, cells eventually develop and select for mutations in specific gene(s) that enable stable and hereditable resistance to stress without physiological costs.^27^ However, beneficial gene mutations can take thousands of generations to develop and become dominant in the population, which necessitates an intermediate adaptation mechanism that enables survival between initial adaptation and heritable resistance. Chromosomal mutations via gene duplication amplification (GDA) events serve as this adaptive intermediate.^1^

GDA events can occur within 100 generations of toxin exposure and involve a 2 to 100 fold amplification of specific genes within the bacterial chromosome.^24^ Furthermore, the kinetics of GDA events are stochastic and copy number can vary significantly within a population.^28^ The size of the amplified region is also highly variable with reported duplicated regions ranging from 18 – 300,000 bp.^8, 24^ Various stressors including carbon source stress, antibiotics, and heavy metals are known to select for high copy number variants, yet the mechanisms by which GDA events are initiated are poorly understood.^25, 29–33^ There is disagreement as to whether GDA events are induced by selective stress, or exist within a population and are selected for as a result of selective pressure.^34, 35^ Additionally, it is typically observed that cells with higher copy numbers (i.e., more amplification events) are more resistant to a given stimulus, however, organism fitness generally decreases as the bacteria maintains these additional copies. Researchers postulate that this genomic redundancy presents the opportunity for beneficial mutations to develop within subsets of the copies without the uncertainty of evolving lethal or non-functional mutations. Thus, duplication protects the genome from deleterious mutations while providing resistance to a given stress to allow for further evolution of resistance genes that become dominant in a population without an associated fitness cost.^36^ ^27^

GDA events occur rapidly to enable survival under in specific stress conditions, but the chromosomal instability of duplication and associated fitness costs have led to duplication events being excluded from discussions of stable mechanisms of bacterial resistance. However, our work demonstrates a stable GDA event in *S. oneidensis* that is triggered by exposure to cobalt and nickel ions released from NMC, enabling stable NMC and metal resistance. More specifically, this gene amplification is uniquely responsive to the combination of cobalt and nickel metal ions. Contrary to other studies evaluating GDA events, there is no clear fitness cost associated with metal-induced amplification in *S. oneidensis,* and, although copy number decreases over time, amplifications (>1) persist for over a thousand generations in the absence of metal stress. No clear compensatory mutations were observed with prolonged metal stress, which further indicates that duplication may serve as a primary, stable, mechanism of multi-metal resistance in *S. oneidensis*. Thus, the work presented herein reaffirms the need to evaluate GDA events as stable mechanisms of bacterial resistance. The importance of understanding the dynamics of GDA events in this metal resistance phenotype is underscored by the growing body of literature linking metal resistance to the dissemination of antibiotics resistance in the environment.

## 2. Results

### 2.1 Gene duplication amplification is a consistent DNA perturbation associated with metal resistance

Previous work from our group has demonstrated that metal ions drive resistance evolution to NMC nanomaterial in *S. oneidensis.* This resistance is stable and hereditable, which implies that chronic metal exposure triggers a genome-level alteration. To explore potential genomic perturbations, metal-resistant cells were interrogated with whole-genome sequencing (WGS). We anticipated a conserved point mutation within the population, however, WGS revealed no consistent point mutations amongst resistant cells (**Table S1**). Rather, WGS resolved an increase in read coverage over a 28,333 bp region of the genome (i.e., coordinate 2,128,908 through 2,157,241) in all metal and NMC-resistant strains. Gene coverage maps indicate that this region of the genome is duplicated between 10 to 25 times after only 48 generations of stress exposure in a minimal media. Notably, no duplication events were observed in WT or passaged control strains (**Figure 1a**). Previous work has demonstrated that metal-ion exposure is sufficient to elicit resistance phenotypes in *S. oneidensis*, and here we demonstrate that resistance to metals and nanomaterials is genetically identical (F**igure 1b**). This further suggests that metal exposure is the primary factor in resistance evolution. Putative annotations of the genes within this region are listed in **Table S2**. WGS results suggest that GDA events are the main driver of NMC and metal resistance over 48 generations, rather than spontaneous point mutations.

**Figure 1:**
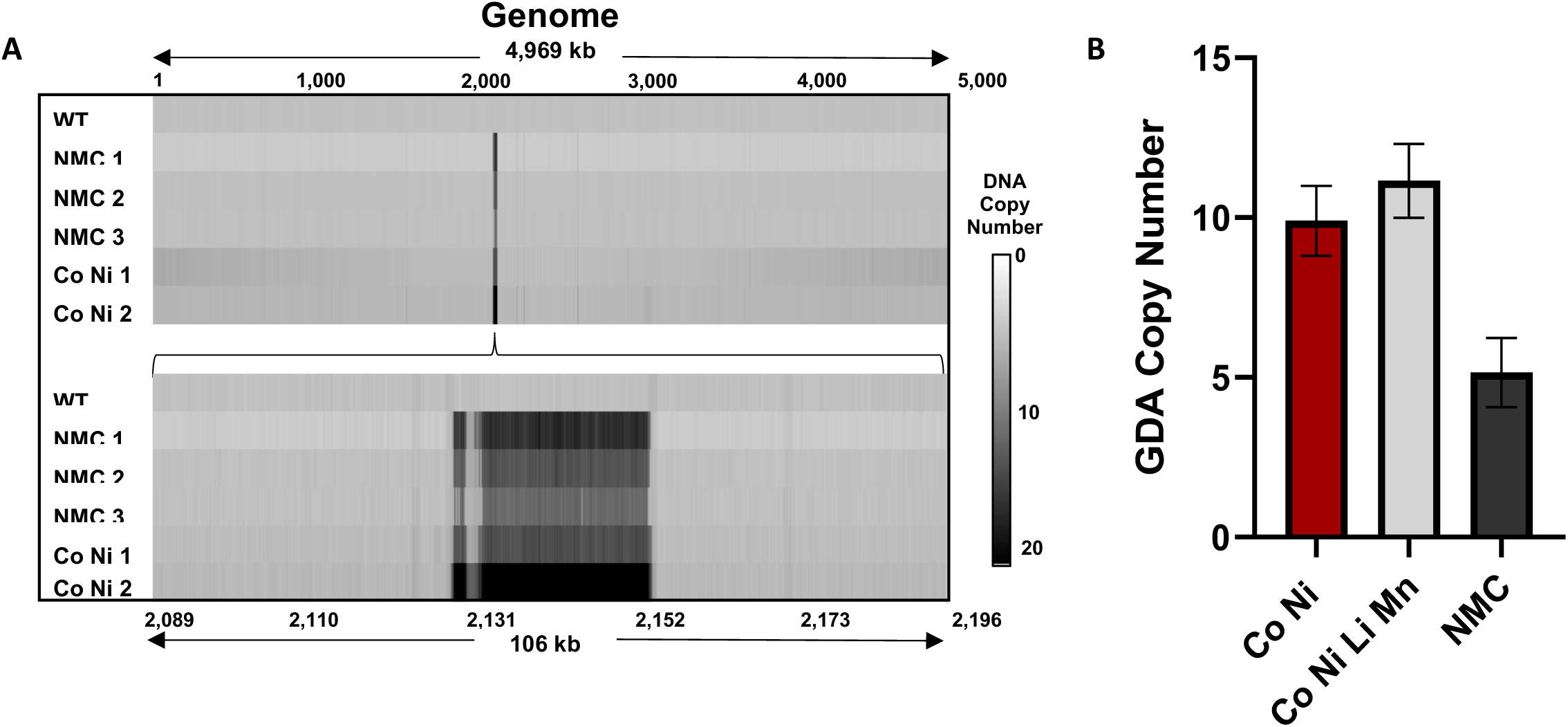
Genomic sequencing and duplication copy number analysis for NMC and metal-resistant strains. A) Whole-genome sequencing read coverage depth of control and NMC-exposed strains. B) qPCR analysis of duplication copy number in cells with evolved resistance to cobalt and nickel, all ions (cobalt, nickel, manganese, and lithium), and NMC. C) qPCR analysis of duplication copy number in cells treated with individual metal ions or a combination of cobalt and nickel.

### 2.2 Cobalt and nickel synergy drive DNA duplication and resistance evolution

Because duplication events are observed in both metal- and nanomaterial-resistant cells, we sought to determine the chemical identify of the stressor(s) that drive gene duplication. Previous work has demonstrated that the combination of cobalt and nickel metal ions most closely mimic NMC toxicity and lithium and manganese impart no additional toxicity to *S. oneidensis*.^37^ As such, we evaluated if concentrations of cobalt and nickel ions equivalent to that released from NMC (25 µM nickel and 10 µM cobalt) in a defined minimal media were sufficient to elicit copy number amplification. Upon exposure to cobalt and nickel for 48 generations, copy number was equivalent to that of cells exposed to the combination of all ions released from the NMC framework (lithium, nickel, manganese, and cobalt) or nanomaterial itself in minimal media (**Figure 1b**). This prompted further investigation into the contributions of cobalt or nickel metal ions, individually. Interestingly, no DNA duplication events were observed with exposure to either 25 µM nickel or 10 µM cobalt monotreatment, whereas approximately eight duplications were observed with combined treatment over this 48-generation timescale (**Figure 2a**). To ensure that duplication is dependent on metal species rather than metal concentration, we exposed *S. oneidensis* to 35 µM cobalt or nickel and similarly observed no duplication event (**Figure S1**). These results indicate that both cobalt and nickel ions are necessary and sufficient to elicit rapid DNA duplication amplification events in *S. oneidensis*.

**Figure 2:**
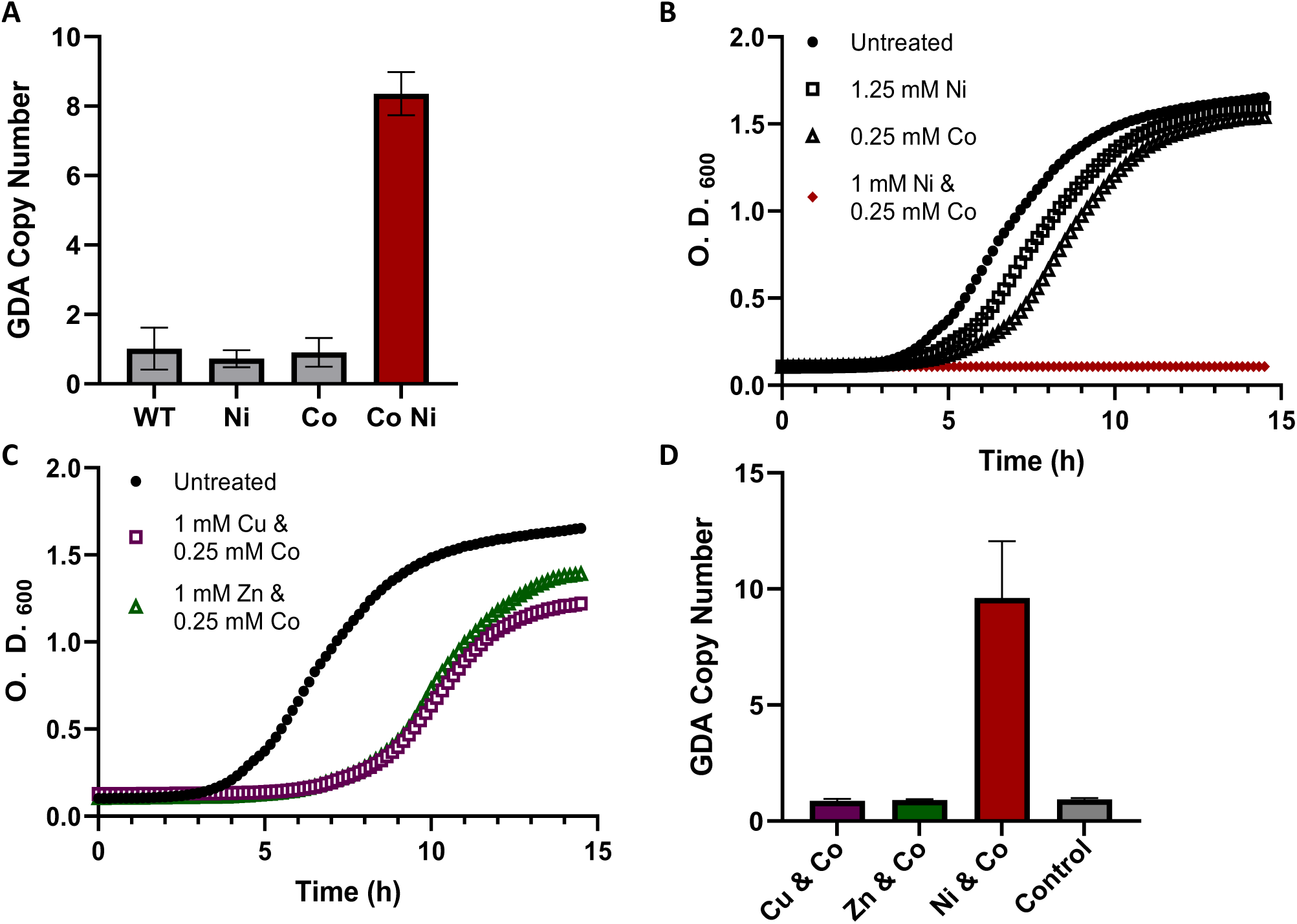
The specific combination of cobalt and nickel interacts synergistically to facilitate gene duplication-mediated resistance in *S. oneidensis*. A) qPCR analysis of DNA duplication copy number upon individual and combination ion exposure. B) Bacterial growth curves with mono and co-treatment of cobalt and nickel metal ions. C) Co-treatment cobalt and either zinc or copper. D) qPCR analysis of DNA duplication copy number after 120 generations with various combinations of metal ions.

To further profile the specific interaction of cobalt and nickel in *S. oneidensis*, we evaluated individual and combined toxicity of cobalt and nickel to WT cells. We sought to determine if the combination of cobalt and nickel was additive or synergistic to better understand how specific metal stress drives GDA. Individual and combined metal MICs in nutrient-rich media were established using a checkerboard assay and synergy was subsequently evaluated by calculating the fractional inhibitory concentration (FIC) value. FIC is used to quantify the change in toxicity between a single compound and multiple compounds to determine if the interaction of the two toxicants is antagonistic, additive, or synergistic. FIC is calculated by comparing the MIC of individual toxins to that of toxins in combination.^38^ The concentration of metal ions used for these assays ranged from 0.25 mM – 4 mM nickel and 0.125 – 2 mM cobalt. The MIC values of nickel and cobalt ions in nutrient-rich media were 4 µM and 2 µM respectively, whereas the MIC for these metals in combination was 1 µM nickel and 0.25 µM cobalt (**Figure 2b**). The calculated FIC value was 0.375, indicating that nickel and cobalt toxicity is synergistic in *S. oneidensis* (**Equation 1**). We postulate that the synergistic interaction of nickel and cobalt facilitates DNA duplication and resistance evolution. To our knowledge, this is the first report of cobalt and nickel synergy in bacterial cells.

### 2.3 Metal synergy and gene duplication amplification is unique to cobalt and nickel

The strong synergistic interaction between cobalt and nickel prompted us to investigate potential synergistic toxicity between other first row transition metals displayed in *S. oneidensis*. The toxicity of copper and zinc metal ions in nutrient-rich media was found to be equivalent to that of nickel (minimum inhibitory concentration equal to 4 µM). Because of this similarity in individual metal ion toxicity, we substituted either copper or zinc for nickel and repeated the previously described checkerboard assays to evaluate potential synergistic toxicity between alternate metals. Interestingly, no novel combination of zinc or copper with cobalt demonstrated synergistic toxicity in *S. oneidensis* (**Figure 2c**). Despite the lack of clear synergy, we exposed cells to the combination of alternate metal ions (i.e., 0.25 mM cobalt and 1 mM of either copper or zinc) for ∼120 generations in LB media and evaluated the potential for elicit gene duplication events. Interestingly, we found that exposure to either copper and cobalt or zinc and cobalt yielded no gene duplications, whereas the combination of nickel and cobalt gave rise to between eight and 13 amplification events (**Figure 2d**). These results indicate that the intracellular interaction between cobalt and nickel is highly specific and DNA duplication events are likely not a general stress response, but instead a coordinated response to specific stimuli.

Interestingly, cells exposed to the combination of copper and cobalt or zinc and cobalt for 120 generations still had a prolonged lag phase indicating that cells did not adapt to metal stress over this time (**Figure S2**). This implies that *S. oneidensis* cannot rapidly adapt to these alternate metal combinations or through the same mechanism that allows for resistance to cobalt and nickel. These results indicate that DNA duplication is a unique response to the combination of cobalt and nickel, and the cell does not have an equivalent duplicative mechanism to deal with toxicity from other combinations of metal ions.

### 2.4 Cells maintain elevated copy numbers and metal resistance in the absence of metal stress

Considering the ecological implications of metal resistance, we sought to determine if elevated copy numbers persist in *S. oneidensis* after an extended recovery period without metal stress. We have previously demonstrated that cells retain metal resistance after culturing for five passages (∼60 generations) in media devoid of metals, which suggests that genetic perturbations persist for multiple generations. However, other work that has looked at gene amplification events demonstrate that amplification events are lost rapidly once stressors are removed. Cell are thought to lose most additional copies within 200 generations without stimuli.

To explore long-term stability of the accumulation of amplification events, we passaged metal-resistant cells (copy number ∼10) into lysogeny broth (LB) without metals for approximately 1,128 generations and evaluated copy number throughout. Interestingly, there was a rapid increase in copy number from ∼6 to 15–20 copies over the first 50 generations. This rapid increase was followed by a steady decrease over the next 350 generations to a copy number of ∼5. After 400 generations, copy number continued to slowly decrease but stabilized at 2–3 copies, and most notably does not return to one copy over the course of our experiment (**Figure 2b**). This data indicates that although copy number decreases, it does not return to one after 1,128 generations in the absence of metals.

Lastly, we evaluated metal tolerance in resistant cells after 1,128 generations with no metal stress. Cell growth was monitored over 20 hours in LB with 1 mM nickel and 0.25 mM cobalt. This concentration of metals is lethal to WT cells, but not to resistant cells. We found that metal-resistant cells that underwent prolonged recovery were able to grow with notably shorter lag phases than WT cells exposed to metals, and growth rate was nearly identical to WT cells with no metal exposure (**Figure 3d**). Together, these data indicate that cells retain metal resistance through prolonged recovery. Importantly, no compensatory mutations unique to metal-resistant cells arose over this recovery period, which suggests that gene duplication remains the primary mechanism of metal resistance in *S. oneidensis* (**Table S3**).

**Figure 3:**
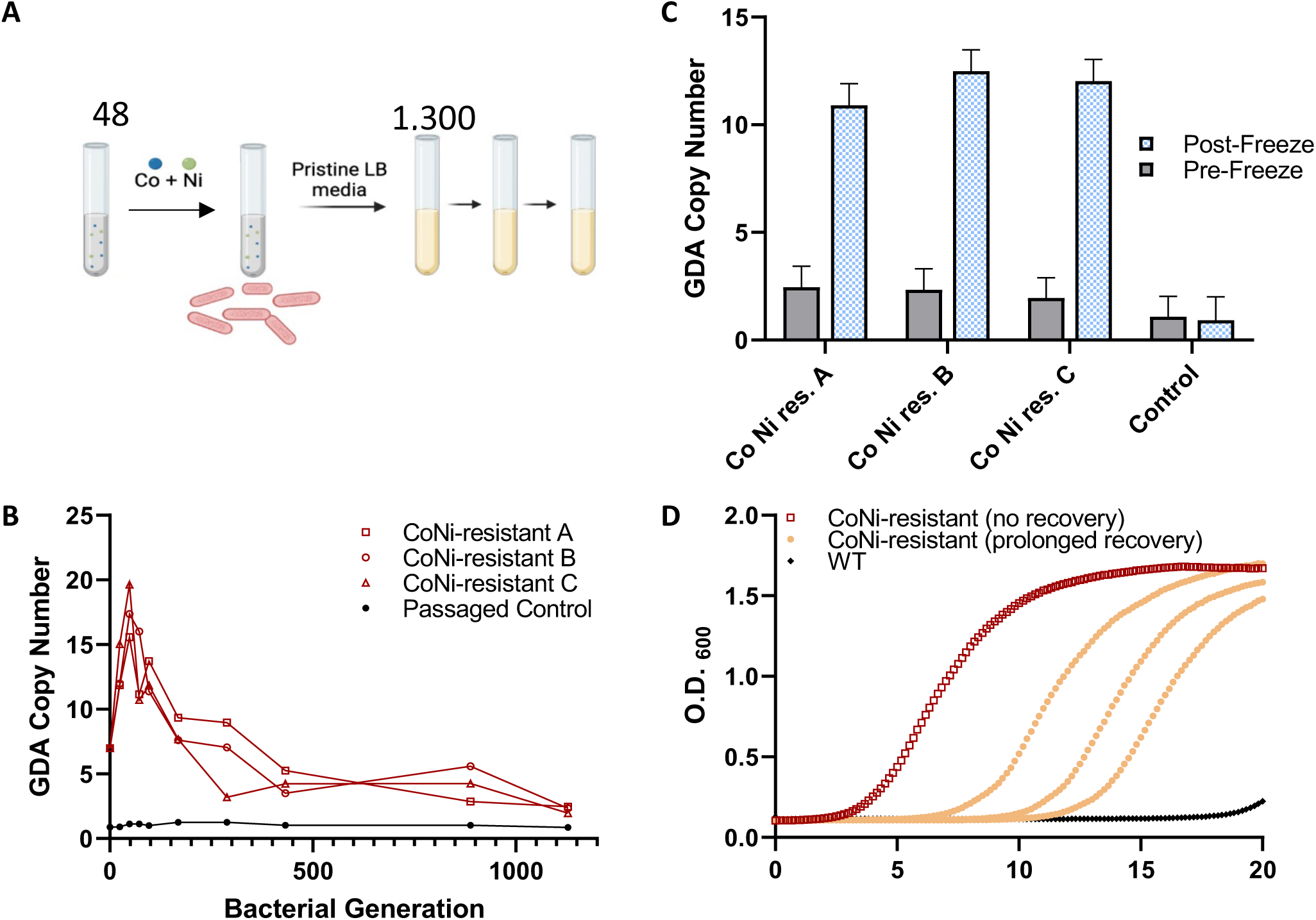
Gene duplication dynamics of metal-resistant cells over 1,128 generations without metal stress. A) Experimental scheme of resistance evolution and cellular recovery. B) qPCR analysis of duplication copy number in three biological replicates over multiple time points throughout the ∼1,100-generation recovery period. C) qPCR analysis of duplication copy number after one freeze-thaw cycle following a 1,128 generations. D) Bacterial growth with metals in cells that have and have not experienced a prolonged recovery period in nutrient rich media.

It was our original intent to use the cells that underwent a prolonged recovery period to study the impact of low copy number on metal resistance. Thus, cells were made into glycerol stocks for future use. Interestingly, when these stocks were streaked on agar plates and grown in pristine media, we found that copy number immediately jumped back up to ∼10 in all metal-resistant replicates (**Figure 2c**). It is important to note that passaged control cells did not have additional copies after cryopreservation. This surprising increase in copy number after cryopreservation highlights the dynamic nature of gene amplification in metal-resistant *S. oneidensis* and the preference cells have for retaining amplification events once they arise.

### 2.5 Metal-induced GDA has no inherent fitness cost

It has been previously reported that each additional kb of DNA results in a 0.15% reduction in cell fitness.^39^ Thus, a 10x amplification event of 28.3 kb in *S. oneidensis* would equate to a 4.25% reduction in fitness. Copy number in metal and NMC-resistant cells is heterogeneous, which suggests that cells have imperfect control over amplification and begs the question of whether there is an observable difference in organism fitness based on copy number. Specific growth rate (µ) and lag phase duration were evaluated in NMC and metal-resistant strains without metal stress. These cells had between 6 and 20 amplification events yet had indistinguishable growth dynamics (**Figure 4a**, **Table 1**). Further growth rate analysis was conducted in metal-resistant cells after a prolonged recovery period, and extended metal exposure (discussed below). Growth dynamics (specific growth rate and lag phase duration) was nearly identical across all strains of *S. oneidensis* regardless of metal exposure or recovery duration (**Figure 4b**, **Table 2**). This further supports our initial observations that copy number does not impact cell fitness under laboratory conditions.

**Figure 4:**
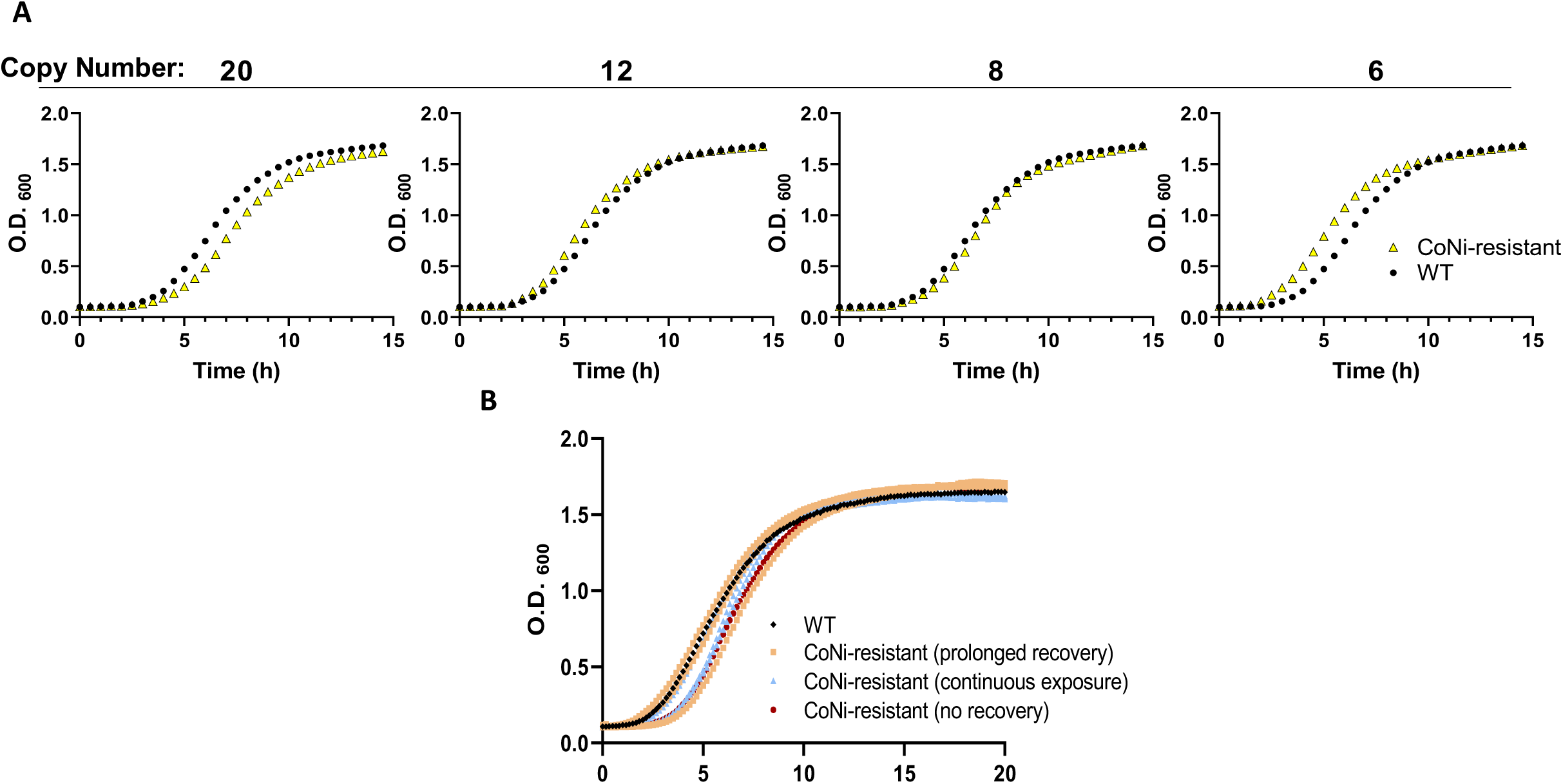
Bacterial growth curves in nutrient-rich media without metal stress. A) NMC-resistant cells with between 6 and 20 duplication events. B) Metal-resistant cells after either 1,128 generations of recovery (CNR_PR), 1,344 generations of continuous metal exposure (PMT), no recover period, or untreated control (WT).

**Table 1.**
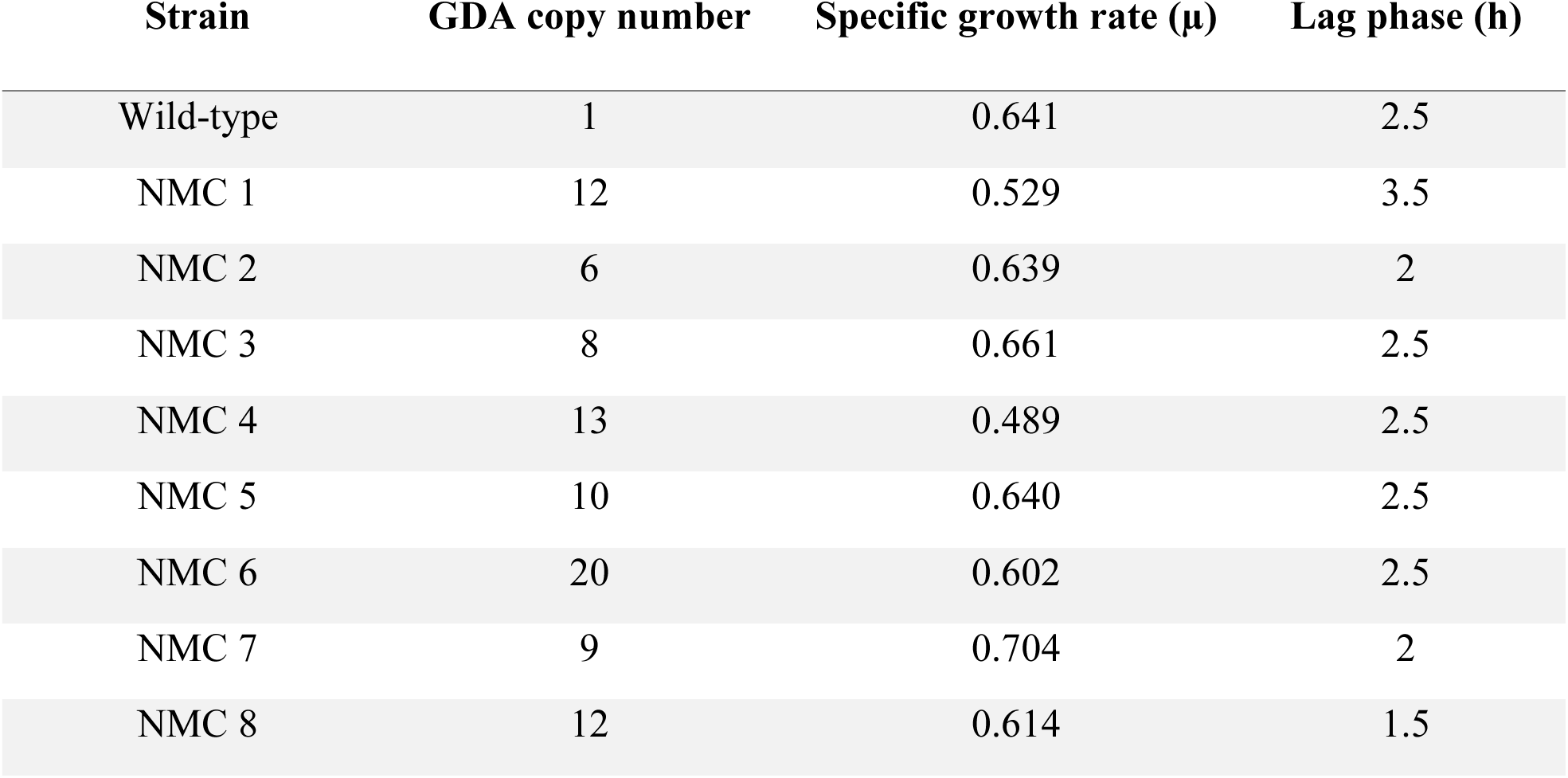
Gene duplication amplification copy number and growth metrics [specific growth rate (µ) and lag phase] in various strains of NMC-resistant *S. oneidensis*.

| Strain | GDA copy number | Specific growth rate ($\mu$) | Lag phase (h) |
| --- | --- | --- | --- |
| Wild-type | 1 | 0.641 | 2.5 |
| NMC 1 | 12 | 0.529 | 3.5 |
| NMC 2 | 6 | 0.639 | 2 |
| NMC 3 | 8 | 0.661 | 2.5 |
| NMC 4 | 13 | 0.489 | 2.5 |
| NMC 5 | 10 | 0.640 | 2.5 |
| NMC 6 | 20 | 0.602 | 2.5 |
| NMC 7 | 9 | 0.704 | 2 |
| NMC 8 | 12 | 0.614 | 1.5 |

**Table 2.** Gene duplication amplification copy number and growth metrics [specific growth rate (µ) and lag phase] in metal-resistant *S. oneidensis* after extended recovery, extended metal exposure.

| Strain | GDA copy number | Specific growth rate ($\mu$) | Lag phase (h) |
| --- | --- | --- | --- |
| Wild-type | 1 | 0.642 | 1.5 |
| Extended exposure 1 | 1.42 | 0.685 | 2.5 |
| Extended exposure 2 | 0.96 | 0.530 | 1.5 |
| Extended exposure 3 | 4.7 | 0.622 | 1.5 |
| Extended recovery 1 | 7.01 | 0.667 | 2.5 |
| Extended recovery 2 | 9.77 | 0.635 | 2.5 |
| Extended recovery 3 | 5.79 | 0.602 | 1.5 |

### 2.6 No clear compensatory mutations develop over 1,000 generations of metal exposure

Because gene duplication amplification events are often described as transient mechanisms of bacterial adaptation, we were interested in probing duplication amplification dynamics and evaluating potential compensatory mutations after extended metal exposure (i.e., > 1,000 generations with metal stress). *S. oneidensis* routinely acquired between 7 and 20 amplification events after 48 generations of metal exposure (25 µM nickel and 10 µM cobalt) in a defined minimal media.^18^ We previously demonstrated that cells have fully evolved metal resistance after 48 generations, meaning that specific growth rates are identical with and without metal stress. Genomic sequencing of these resistant populations revealed between 10 and 25 amplification events (discussed previously). Considering the variability in amplification frequency, we were interested in the dynamics of amplification over thousands of generations of metal exposure. Because growth rate of *S. oneidensis* in minimal media is slow (∼6 hour doubling time), metal exposure was adapted for nutrient-rich LB, (∼30 minute doubling time).

We first optimized metal concentration to find a similar toxicity in LB as previously observed in minimal media. It was found that 0.25 mM cobalt and 1 mM nickel was lethal to *S. oneidensis* in LB, but chronic exposure led to resistance evolution and the onset of duplication events. Cells were then passaged for approximately 1,344 generations in LB with metal stress and copy number evaluated at various timepoints. Copy number increased rapidly over the first 72 generations from 1 to ∼10. After 72 generations, duplication events continued to accumulate before reaching a maximum copy number of ∼15 by generation 500. Copy number ultimately decreased and equilibrated around 10 after an additional 700 generations of metal exposure (**Figure 5b**). These results demonstrate that copy number accumulation is dynamic, but cells retain 10+ copies upon prolonged metal stress.

**Figure 5:**
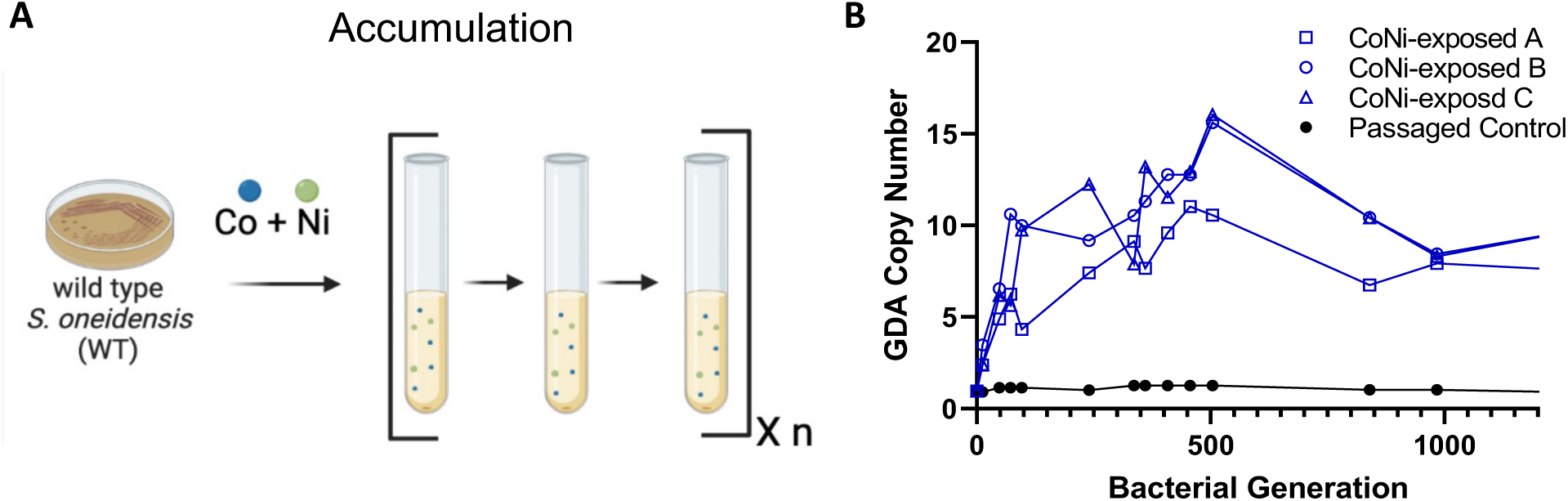
Copy number accumulation over extended duration of metal exposure. A) Experimental scheme. B) qPCR analysis of copy number at various timepoints throughout 1,344 generations of exposure to the combination of 1 mM nickel and 0.25 mM cobalt in three strains of *S. oneidensis* and a parallel passaged control.

Previous analysis of gene duplication amplification events demonstrate that cells acquire compensatory point mutations after 100 generations of continual stress.^27^ These point mutations became the dominant mechanism of resistance and unnecessary copies are lost. To investigate whether 1,344 generations of metal treatment led to the evolution of compensatory point mutations, we submitted cells that underwent 1,344 generations of metal treatment for WGS. Interestingly, there were no consistent point mutations among sequenced strains, and one strain displayed no additional genomic alterations upon extended metal exposure (**Table S3**). The lack of evolution of compensatory mutation is supported by evidence that cells retained ∼10 amplifications after 1,344 generations of metal treatment and that all strains have similar fitness to untreated *S. oneidensis* in the absence of metal stress (**Figure 3b**). Taken together, these observations suggest that gene duplication amplification is likely the primary mechanism of metal resistance in *S. oneidensis*.

### 2.7 Putative cation efflux pump is necessary for cobalt and nickel resistance evolution

Putative annotations of the genes within the duplicated region indicate that one of the duplicated genes is a cation diffusion facilitator (CDF) family protein, locus tag SO_2045. Because efflux proteins have a well-documented role in metal homeostasis, we were particularly interested in the role of this protein in our system.^40–42^ Phylogenetic predictions of SO_2045 suggest that it is selective to cobalt and nickel metal ions.^43^ Thus, we sought to determine the role of SO_2045 in cobalt and nickel resistance evolution by using a SO_2045 transposon knockout strain obtained from Dr. Buz Barstow (denoted SO_2045::TnKan).^44, 45^ Optical density measurements of WT and SO_2045::TnKan were used to evaluate differences in metal toxicity between the two strains. Cobalt mono-treatment (0.25 mM) in LB media was notably more toxic in the efflux mutant when compared to WT (**Figure 6a**). However, sensitivity to 1 mM nickel remained similar between the mutant and WT strains, which suggests that the efflux pump is critical for cobalt tolerance, but nickel tolerance is under the control of an alternate system. Further transcript analysis shows that SO_2045 is ∼100 fold upregulated in metal-treated cells over 48 generations in minimal media relative to passaged controls, further confirming the importance of the gene (**Table S4**).

**Figure 6:**
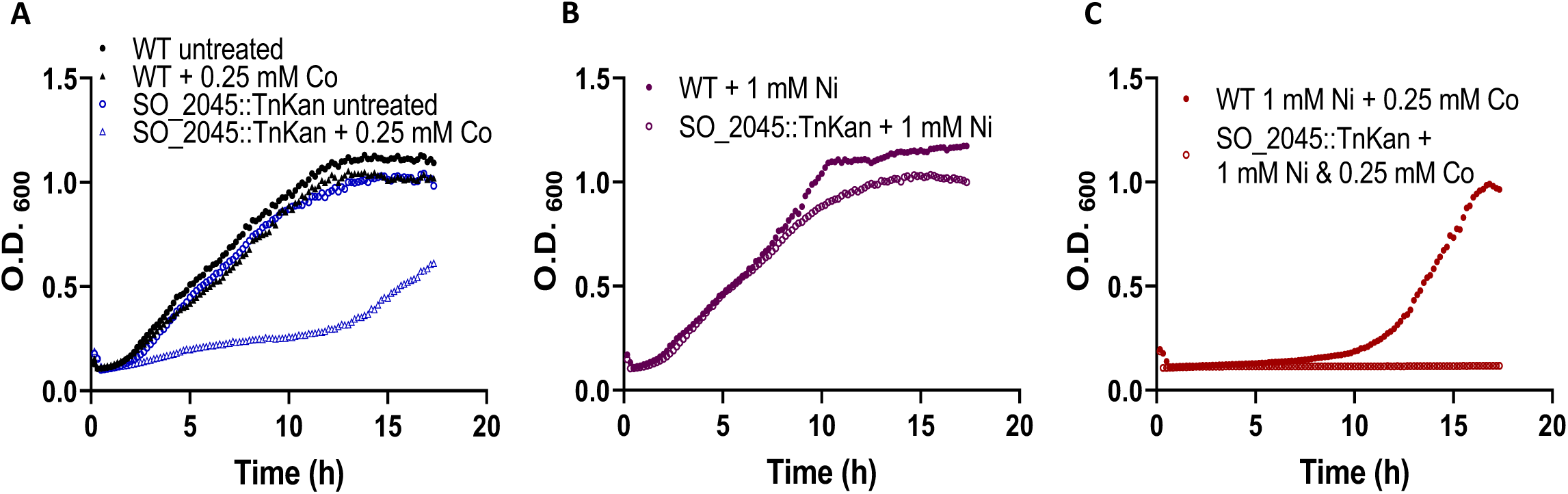
Cobalt and nickel toxicity in CDF-efflux mutant (SO_2045::TnKan). Bacterial growth with A) 0.25 mM and 0.5 mM cobalt, B) 1 mM nickel, and C) the combination of 1 mM nickel and 0.25 mM cobalt in WT and SO_2045::TnKan.

We next evaluated the effect of combined cobalt and nickel treatment in the efflux deletion strain. Growth data reveals that the combination of metal ions is lethal in ΔSO_2045 relative to WT cells (**Figure 6b**). These data demonstrate that SO_2045 has a critical role in *S. oneidensis* adaptation to cobalt toxicity. However, this CDF family efflux protein does not provide insight on the mechanism of cobalt and nickel synergy or why this metal combination specifically induces GDA events. Although we have shown that SO_2045 has a critical role in metal resistance evolution, future work will involve further analysis of genes beyond the duplicated region to more completely assess the mechanism of cobalt and nickel synergy in *S. oneidensis*.

## 3. Discussion

In this study, we identify a consistent gene duplication amplification event that arises in response to chronic exposure to cobalt and nickel in *S. oneidensis*. A putative CDF-family efflux pump within the 28-kb duplicated region was found to mediate cobalt toxicity, but the intracellular mechanism that drives synergy between cobalt and nickel ions remains unknown. Evidence suggests that CDF-family efflux proteins evolved through gene duplication, and are therefore likely to be associated with mobile genetic elements (MGE).^46^ However, duplication of CDF genes has not previously been reported as a mechanism of metal-resistance in prokaryotes. We describe a highly dynamic metal-induced duplication amplification event in *S. oneidensis* that evolves rapidly, persists for over 1,000 generations in the absence of selection pressure, and can be re-stimulated with stress that does not trigger initial duplication amplification events. Furthermore, in contrast to previous studies that look at gene duplication, we observe that duplication events do not cause fitness defects in metal-resistant cells. This lack of fitness cost indicates that gene duplications may persist within a microbial community longer than previously thought due to chronic stress. This suggests that gene duplication amplification events should be more widely considered as a mechanism of bacterial resistance to environmental toxins and antibiotics.

Chronic exposure to toxins can lead to resistance, which includes the evolution of and selection for beneficial genetic mutations in specific cellular components.^47^ However, mutations in individual genes evolve slowly and thus, gene duplication events have been observed as an intermediate, ephemeral response to toxins. The transient nature of duplication events is presumed to contribute to heteroresistance, where a subset of the bacterial population is significantly less susceptible to a toxin, but the population does not retain resistance if direct stress is removed.^28, 48^ Most research indicates that additional copies of genes are rapidly lost in the population in the absence of selection pressure due to the high fitness cost associated with harboring additional DNA. The loss of duplication events leads to the resensitization of an organism to a given toxin if the organism has not developed a stable, compensatory mutation. This loss of duplication events within a population drives the transient nature of duplication-mediated resistance. However, the stability of the DNA duplication amplification events presented in this work suggests that duplication events may serve as a primary resistance mechanism in some systems and can exist without fitness costs.

Prolonged culturing of metal-resistant cells in the absence of metal stress led to a decrease in the number of amplification events per organism, but in no cases resulted in the complete loss of duplication. It is possible that duplication events might be eliminated from the population if cells were cultured without stress for even longer durations, however, considering that routine cryostress can uniquely elicit additional amplification events in metal-resistant cells, it is unlikely that cells will be able to obtain this prolonged recovery period in a complex environmental matrix. Similarly, organisms maintained ∼10 copies throughout prolonged metal exposures even in strains that acquired additional point mutations over the course of treatment. In addition to possessing similar duplication amplification dynamics, strains that developed point mutations had no clear fitness advantage or increased metal resistance compared to strains without mutations. Because cells that acquire additional point mutations do not harbor a competitive advantage over cells that have no additional mutations, we postulate that gene duplications are the primary driver of the resistance phenotype.

We demonstrate that this GDA event arises as a specific response to cobalt and nickel bimetal stress. To the best of our knowledge, this is the first demonstration of metals interacting synergistically to initiate GDA events despite the many studies that report bimetallic nanoparticles or dual metal-ion treatments leading to enhanced bacterial inhibition relative to single-metal treatments.^49–51^ The mechanisms of metal ion synergy remain largely elusive, but it is hypothesized that increased toxicity is due to dual inhibition of intracellular processes such as DNA damage, membrane integrity, metabolic function, or antioxidant depletion.^51, 52^ Considering these mechanisms of toxicity, it is not clear why the combination of cobalt and nickel synergizes while the combination of cobalt and copper or zinc does not. Furthermore, current understanding of mechanisms of metal resistance is largely limited to general stress responses (i.e. efflux, sequestration, and enzymatic conversion).^53^ Reversible GDA events in known metal-resistance genes have been observed as a temporary response to individual nickel, copper, or cadmium metal ion stress, but these studies do not address the potential impact of bimetallic stress.^54–56^ Thus, the intracellular mechanism that elicits GDA events as a result of specific cobalt and nickel stress in *S. oneidensis* requires further investigation and is of growing importance due to the increasing abundance of cobalt and nickel in the environment.^57, 58^

To understand the ecological impact of the evolution of metal resistance in *S. oneidensis*, we must consider the potential for resistance to be shared amongst other organisms. MGEs are critical for horizontal gene transfer (A) between bacterial species.^59–61^ Insertion sequences and transposons can move to new sites on the chromosome or to plasmids within the cell.^62^ This mobilization is facilitated by transposase enzymes, and thus, the association of resistance genes with transposases and insertion sequences is a critical step in HGT.^63–65^ Putative functional annotations of the genes duplicated in metal-resistant *S. oneidensis* reveal multiple transposases and integrases that flank approximately 17 cargo genes. The structure of this genomic region and association of cargo genes with recombinase enzymes indicates that this MGE may ultimately be exported from *S. oneidensis* and transferred to other organisms in the same environmental niche.^64, 66, 67^ Furthermore, the MGE may be transferred to a pathogenic species, which is particularly concerning in the context of the growing antibiotic resistance crisis.^68–70^ Alternatively, the amplified genes may trigger a defense response and lead to the production of novel toxins, which could have deleterious effects on the broader microbial community.^71^ Further work is needed to elucidate the inter-organism transfer of the MGE studied herein, but is important to pursue to understand the contribution of GDA towards resistance evolution.

The evolution and dissemination of genes that facilitate resistance to environmental toxins is of interest because environmental bacteria and aquatic environments are known to be reservoirs of antibiotic resistance genes.^59, 72–75^ Indeed, certain antibiotic resistance genes are known to originate from *Shewanella* species.^76–78^ *S. oneidensis* exists in the microbiome of some aquatic species and co-exist with known human pathogens, such as *Vibrio cholerae*, which increases the probability that genes from *S. oneidensis* may be transferred to a pathogen.^79^ Putative functions of other protein-coding genes within the MGE may provide additional fitness benefits to organisms, or protection from environmental toxins not identified in this study. Furthermore, certain genes may be beneficial for survival under antibiotic stress, which underscores the importance of studying gene duplication. Additional analysis of the function of genes duplicated in metal-resistant *S. oneidensis* is required to fully categorize the physiological benefits associated with gene duplication and the impact of sharing these genes with other organisms.

Beyond facilitating cell survival under stress conditions, gene duplication facilitates further genomic adaptation through the evolution of novel resistance genes.^80–83^ Redundancies in the genome provide opportunity for novel mutations to develop without threatening bacterial viability. Thus, the stable GDA event presented here may provide a platform for novel resistance to other environmental stressors or antibiotics to arise in *S. oneidensis* and other organisms.

It is known that gene duplications arise in response to external stress and a diverse array of stimuli can elicit duplication events. However, the mechanism through which specific stimuli elicit particular duplication events is unknown due to the complexity of intracellular signaling that leads to duplication. Here, we demonstrate that the specific combination of cobalt and nickel is required to elicit duplication events in *S. oneidensis*. This suggests that these metal ions have unique intracellular targets that interact to ultimately trigger duplication amplification events. Additional work is needed to elucidate the specific mechanism through which these metal ions elicit duplications.

Future studies will seek to address the ability of genes within the duplicated region to be spread to other organisms along with developing a more complete understanding of how cobalt and nickel facilitate duplication events and changes in biophysical properties brought on by resistance. Together, the work presented herein suggests that GDA is a more stable mechanism of bacterial resistance than has previously been appreciated. Thus, duplication amplification events should be characterized and considered more thoroughly among resistant populations as these genomic perturbations may provide an avenue for novel mutations and facilitate the spread of novel antibiotic resistance genes.

## Acknowledgements

This work was supported by the National Science Foundation under Grant No. CHE-2001611, the NSF Center For Sustainable Nanotechnology. The NSF CSN is part of the Center for Chemical Innovation Program. A. Gavin acknowledges support from the Lester C. and Jan M. Krogh Excellence Fellowship. The authors gratefully acknowledge Cameron Kitzinger and Dr. Buz Barstow for sharing SO_2045::TnKan from their whole-genome transposon insertion library.

## Materials and Methods

### DNA extraction & whole genome sequencing

Single colonies were picked from luria broth (LB) agar plates and inoculated into 5 mL of LB Lennox broth. Cultures were grown overnight, diluted to an OD_600_ of 1.0. and 1.8 mL was used for DNA extraction with a Qiagen DNeasu UltraClean Microbial Kit according to manufacturer’s instructions. DNA was eluted in RNase-free water and measured for concentration and purity using a NanoQuant Plate and Tecan Spark Plate reader. DNA was then either submitted for sequencing by SeqCenter (Pittsburgh, PA) or prepared for qPCR copy number analysis.

### Whole-genome assembly and copy number analysis

Sequencing data was returned as two fastq files per strain corresponding to forward and reverse reads. Reads were mapped and aligned to a reference genome assembly for *Shewanella oneidensis* (GCF_000146165.2). Mapping was done with BWA to produce a .bam file and was subsequently converted to a bigwig file using bamCoverage. All alignment and file conversion was performed via Galaxy (version 24.2). Bigwig files were visualized using integrated genomics viewer (IGV).

### GDA accumulation

*Shewanella oneidensis* MR-1 (ATCC BAA1096) cells streaked on lysogeny broth (LB) agar plates at 30 °C. Three colonies were picked and inoculated in 5 mL LB media each for 12 hours after which cells were sub-cultured in technical triplicate. Cells were subsequently passaged every 24 generations (∼12 hours) and aliquots were collected intermittently for DNA extraction at and copy number analysis.

### GDA loss

Three metal-resistant strains were streaked on LB agar plates. Three colonies were picked from each strain and grown in 5 mL pristine LB media. Unperturbed *S. oneidensis* has a doubling time of ∼30 minutes in LB media, thus, there are ∼24 generations in a 12-hour passage. Cells were diluted 1:250 and passaged every 12 hours for 25 days, a total of 1,200 generations (50 passages) in pristine media. Aliquots were taken intermittently for DNA extraction and copy number analysis.

### Determination of transposon copy number by real-time qPCR

DNA concentration was diluted to 10 ng/µL for copy number evaluation. Relative copy number was determined using a qPCR-based method.^84^ SO_4604 (putative annotation, *sulA*) was selected as the reference gene because there is only one copy in the genome, and it lies outside the duplicated region, which means that there should only remain one copy even after duplication events occur. SO_2041 (putative annotation, *rmuC*) was chosen as the target gene. SO_2041 lies within the duplicated region and should thus have a detectable fold change that corresponds to TE copy number with duplication events. All copy number assessments were normalized to DNA extracted from the same unpassaged control samples.

### Metal MIC and synergy

1M stocks of ZnCl_2_ CuCl_2_ CoCl_2_ NiCl_2_ were prepared in MiliQ H_2_O and sterile filtered. Metal stocks were diluted to appropriate concentrations as follows: 1 mM Ni, Cu, or Zn and 0.25 mM Co. Growth curves were measured in a 96-well plate (in a Tecan Spark microplate reader). Optical density measurements at 600 nm were taken every 10 minutes. MIC is defined as treatments that yield an optical density less than 0.2 after 20 hours of growth in LB media.

**Equation 1.** Fractional inhibitory calculation to determine synergy between cobalt and nickel metal ions.

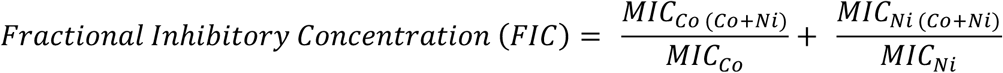

