## Supporting Information Document for "Cobalt and nickel ion synergy promotes gene duplication amplification and enables stable metal resistance in *Shewanella oneidensis*"

**List of Tables**

**Table S1.** Mutations in metal- and NMC-resistant *S. oneidensis* identified through whole-genome sequencing.

**Table S2.** Putative annotations of duplicated genes.

**Table S3.** Mutations identified after prolonged metal exposure in *S. oneidensis*.

**Table S4.** Differences in SO_2045 transcription after 48 generations of metal exposure.

**List of Figures**

**Figure S1.** Copy number analysis upon 35 μM single-ion metal treatment.

**Figure S2.** Bacterial growth after 120 generations of metal exposure.

**Table S1.** Mutations identified in whole-genome sequencing of metal- and nanomaterial-resistant *S. oneidensis.*

| Strain | Location | Locus | Original | Mutation | Gene |
|---|---|---|---|---|---|
| Ion-resistant | Plasmid | 38,067 | ATTTTTTTAT | ATTTTTTTTATG, insertion | SO_A0176, hypothetical |
| NMC-resistant | Chromosome | 256,847 | G | C | *rpsK*, SO_0254 |
| NMC-resistant | Chromosome | 257,910 | T | C | *rpoA*, SO_0256 |
| NMC-resistant | Chromosome | 259, 251 | G | A | Non-coding |
| NMC-resistant | Chromosome | 865,439 | | TCAGCAT insertion | Non-coding |
| NMC-resistant | Chromosome | 1,487,743 | G | A | Non-coding |
| NMC-resistant | Chromosome | 3,388,411 | | GTCATACGGTAGAG deletion | *flgB*, SO_3250 |
| NMC-resistant | Chromosome | 4,902,469 | C | A | SO_4700 |

**Table S2.** Putative annotations of genes identified within the duplicated region in metal- and nanomaterial-resistant *S. oneidensis*.

| **Locus Tag** | **Gene** | **Putative Annotation** | **Cellular location** |
|---|---|---|---|
| SO_2032 | tnpA | ISSod1 transposase | Cytosol |
| SO_2034 | | Hypothetical protein | |
| SO_2035 | tnpA | ISSod25 transposase | Inner membrane |
| SO_2036 | | ISSod25 integrase Int_ISSod25 | cytosol |
| SO_2037 | intl | Integron integrase | cytosol |
| SO_2039 | | Signaling protein with EAL domain | Inner membrane |
| SO_2040 | sltY | Soluble lytic murein transglycosylase | periplasm |
| SO_2041 | rmuC | DNA recombination protein | cytosol |
| SO_2042 | yedY | Oxidoreductase molybdenum-binding subunit | |
| SO_2043 | yedZ | Oxidoreductase cytochrome b subunit | Inner membrane |
| SO_2044 | gloA | Lactoylglutathione lyase | cytosol |
| SO_2045 | | Cation efflux protein CDF family | Inner membrane |
| SO_2046 | | Transcriptional regulator MarR family | cytosol |
| SO_2047 | | Prolyl oligopeptidase family S9 protein | periplasm |
| SO_2048 | | Phosphoethanolamine transferase | Inner membrane |
| SO_2049 | | Diguanylate cyclase | cytosol |
| SO_2050 | | Methylase | cytosol |
| SO_2051 | | Hypothetical protein | |
| SO_2052 | | FAD-dependent oxidoreductase | cytosol |
| SO_2053 | | Transcriptional regulator LysR family | cytosol |
| SO_2054 | frmA | S-(hydroxymethyl)glutathione dehydrogenase | cytosol |
| SO_2055 | frmB | S-formylglutathione hydrolase | |
| SO_2057 | tnpA | ISSod1 transposase | |

**Table S3.** Mutations identified in cells that experienced prolonged metal treatment.

| Locus | CNR | Prolonged Metal Exposure1 | Prolonged Metal Exposure 2 | Prolonged Metal Exposure 3 |
|---|---|---|---|---|
| 257910 | X | X | X | X |
| 650,960 | X | CRP transcriptional regulator | X | X |
| 1,162,400 | X | intergenic peptidase | X | X |
| 1,386,228 | X | X | X | SO_1329 |
| 1,699,961 | X | X | X | X |
| 2,047,497 | X | X | X | SO_1944 |
| 2,689,107 | X | X | X | X |
| 2,832,134 | X | X | X | X |

**Table S4.** Transcript analysis of SO_2045 after 48 generations of metal-treatment. $Log_2$ fold change relative to passaged control.

| Gene | $Log_2$ Fold Change | Adjusted p value |
|---|---|---|
| SO_2045 | 6.01714 | 2.02 E-55 |

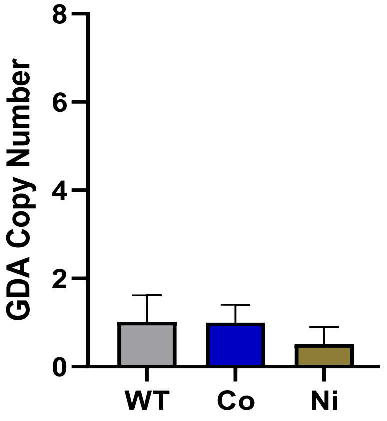


**Figure S1.** Copy number analysis after 48 generations of 35 µM single-ion metal treatment compared to passaged control (WT).

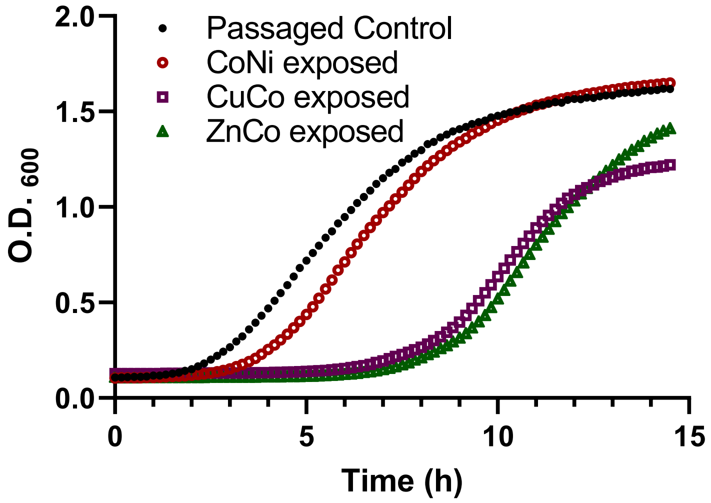


**Figure S2.** Bacterial growth analysis after 120 generations of exposure to either Cu & Co, Zn & Co, or Ni & Co.
